# PlasmidLM: A Promptable DNA Language Model via Verifiable-Reward Post-Training

**DOI:** 10.64898/2026.05.19.725242

**Authors:** McClain Thiel, Chris P. Barnes

## Abstract

Generative DNA models are typically next-token completers: they extend a sequence but offer no native interface for telling the model what to make. PlasmidLM is a promptable DNA language model for plasmids. A designer supplies a human-readable component specification, for example a high-copy *E. coli* vector with kanamycin resistance and an EGFP reporter, and the model generates the corresponding multi-kilobase construct in a single autoregressive pass. Prompts are unordered sets of named-part tokens at the granularity of biological shorthand, not learned latent codes or rigid grammars. We evaluate outputs along two axes: a sequence is *viable* if structurally plausible as a plasmid, and *faithful* if its detected components match the prompt. Their conjunction is the *useful-plasmid rate*, the primary metric we report. On a held-out 1,000-prompt benchmark, the post-trained model achieves a useful-plasmid rate of 48.5% at single-shot decoding and 89.7% under best-of-4 sampling. Verifiable-reward post-training with GRPO against a 660-entry sequence motif registry improves the useful-plasmid rate across all sampling budgets. We release the 19.3M-parameter model, evaluation suite, and a paired benchmark of prompt-sequence pairs.

## 1 Introduction

Designable DNA is the task of taking a high-level specification of a desired construct and producing a sequence that realises it. It is the daily work of synthetic biology, from cloning and antigen design to gene-therapy vector assembly and CRISPR guide-vector pipelines. Genomic language models do not natively support it. Nucleotide Transformer [Dalla-Torre et al., 2025], DNABERT-2 [Zhou et al., 2024], and HyenaDNA [Nguyen et al., 2023] are pretrained for discriminative downstream tasks (variant-effect prediction, regulatory annotation) and expose no generative interface. Evo 2 [Brixi et al., 2026] is the most ambitious recent generative effort: it operates at genome scale and accepts a small catalogue of species tags as prefix conditioning, but is otherwise a sequence completer. A handful of conditional generators target short regulatory elements [Sample et al., 2019, Stark et al., 2024, Taskiran et al., 2024], but at the level of an entire multi-kilobase construct we are not aware of any prior DNA model that accepts a human-readable component specification and returns a candidate sequence intended to satisfy it.

This task calls for a different evaluation than completion. Perplexity scores how well a model continues a sequence; it does not score whether the sequence satisfies a specification. We separate two axes. A sequence is *viable* if it is structurally plausible as a plasmid: a full complement of essential elements arranged in a configuration compatible with replication and expression. A sequence is *faithful* if its detected components match the components requested in the prompt. Their conjunction is the *useful-plasmid rate*, the primary metric we report. Faithfulness is currently absent from the genomic-LM literature because there are no standard DNA models with prompts to be faithful to.

Plasmids are a deliberate testbed for this question. They are compact (typically 4 to 10 kb), modular, deposited at scale with lab-supplied metadata, and amenable to fully automated annotation: an existing pipeline (pLannotate [McGuffie and Barrick, 2021]) can identify origins, selection markers, regulatory elements, and reporters in a candidate sequence and return a structured component list. The Addgene repository contributes over 100,000 sequenced plasmids deposited by working laboratories, each carrying lab-supplied metadata about its intended use. Crucially, the same pipeline that scores faithfulness at evaluation time supplies the verifiable reward used during post-training, so the model is reinforced for the property that is being measured.

This paper makes three contributions. *Promptable DNA generation:* PlasmidLM is a 19.3M-parameter plasmid language model that accepts a human-readable component specification as a prompt and generates a candidate multi-kilobase construct in a single autoregressive pass. *An evaluation framework for promptable DNA:* the viable/faithful decomposition and the useful-plasmid rate as a summary metric, grounded in a 660-entry motif registry that ties human-readable token names to concrete sequence-level references. *Verifiable-reward post-training for DNA:* GRPO against the same registry used for evaluation, which improves the useful-plasmid rate across single-shot and best-of-*K* decoding. The post-trained model produces useful plasmids in 48.5% of single-shot generations and in 89.7% of best-of-4 generations on a held-out 1,000-prompt benchmark. Figure 1 shows three generations spanning the quality range to calibrate what the aggregate numbers mean concretely.

**Figure 1:**
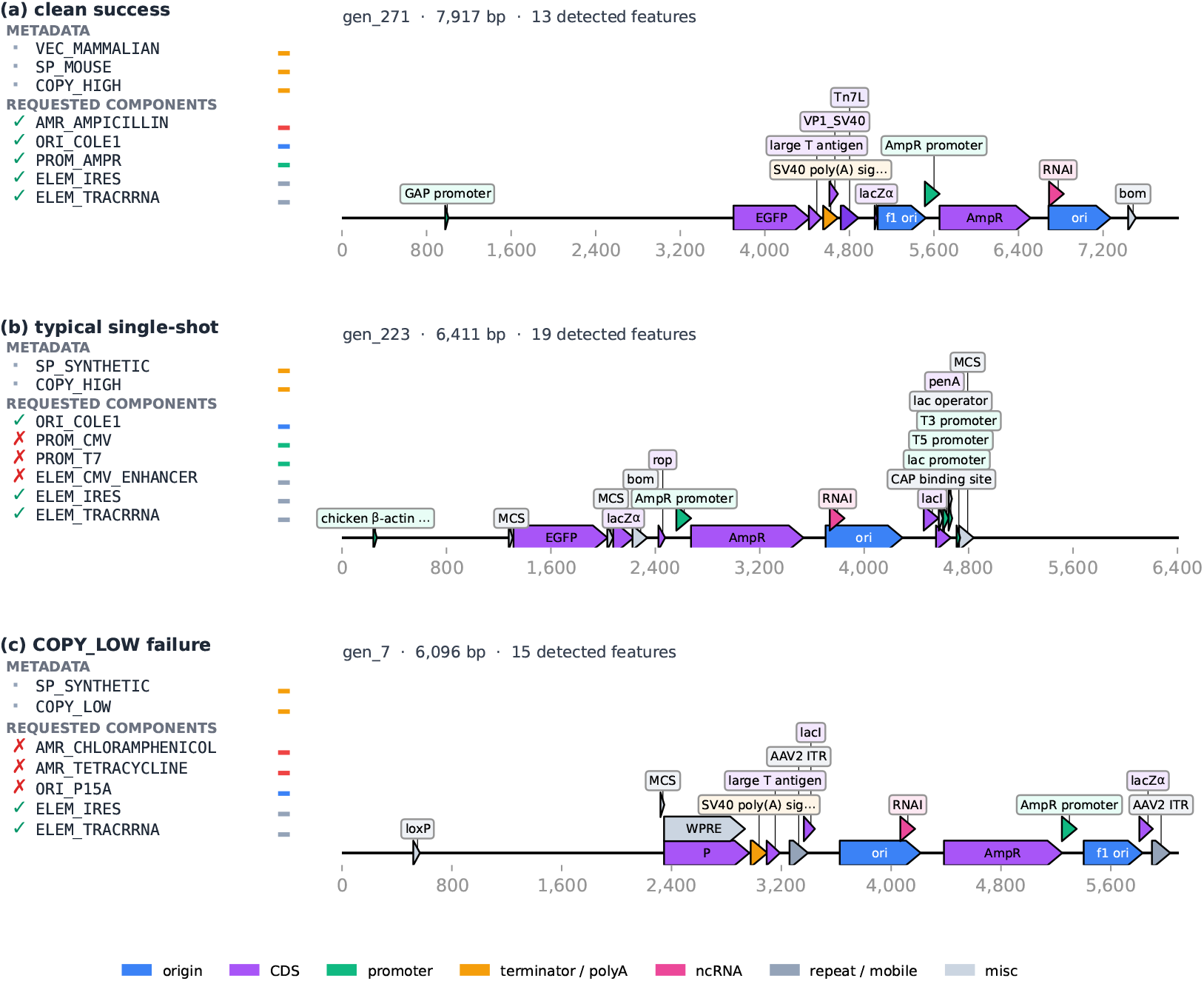
Exemplar generations spanning the quality range. Each row: prompt (left) and pLannotate annotation map (right). (a) Clean success: 5/5 tokens recovered, coherent mammalian/*E. coli* shuttle vector. (b) Typical single-shot: 3/6 tokens recovered, plausible plasmid on a different backbone. (c) `COPY_LOW` + rare-AMR failure: model emitted high-copy origins and ampicillin resistance instead of the requested low-copy + chloramphenicol/tetracycline.

**Figure 2:**
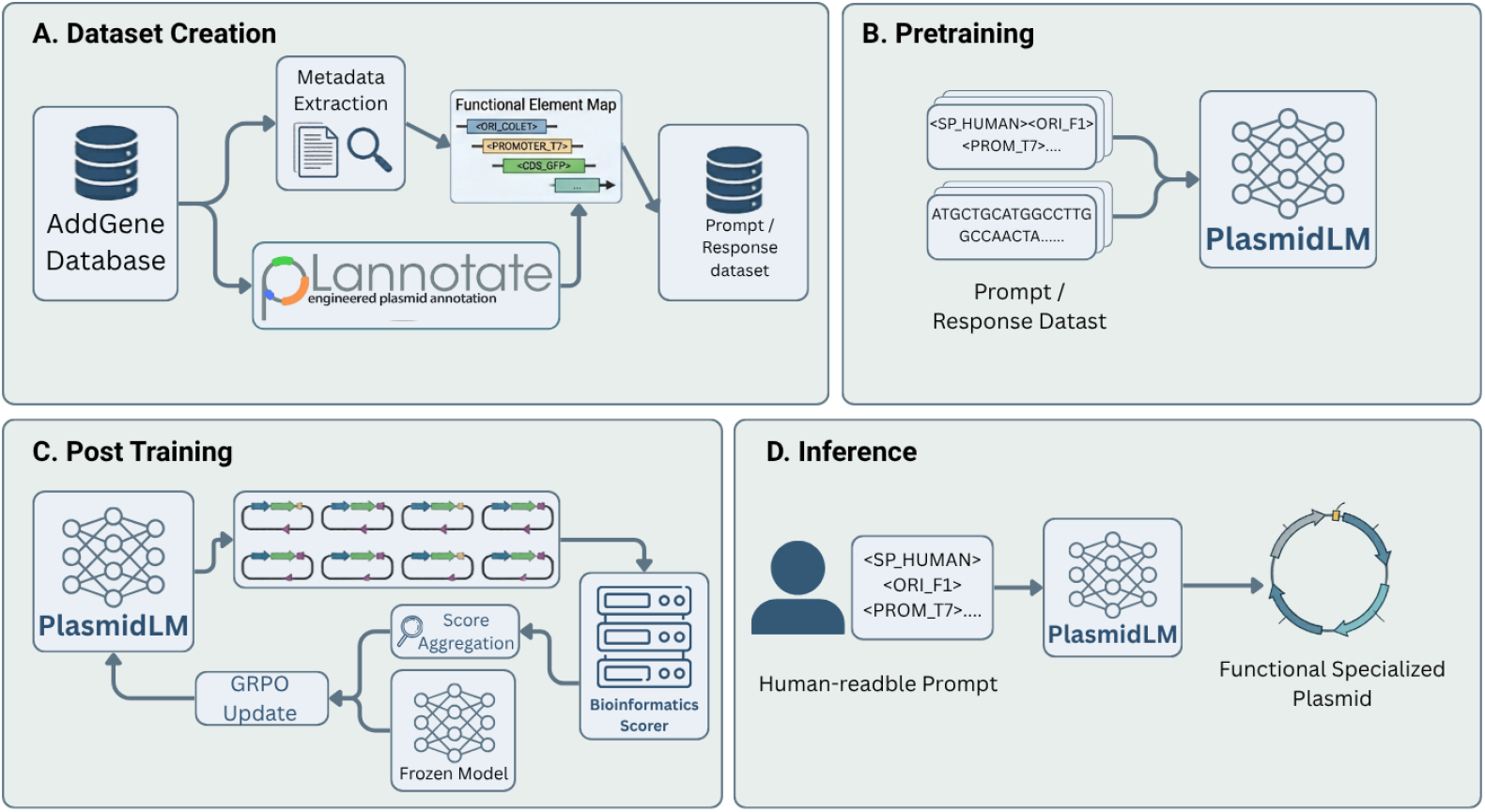
PlasmidLM pipeline. **(A)** Addgene sequences annotated by pLannotate; functional elements mapped to hard tokens via the motif registry, supplemented by soft metadata tokens. **(B)** Prompt-paired pretraining with next-token prediction. **(C)** GRPO post-training: *G*=8 completions scored by BLAST against the registry; group-normalised rewards with KL constraint to the frozen reference. **(D)** Inference: user-supplied token list conditions autoregressive generation.

## 2 Related work

### 2.1 Generative DNA models

Genomic language modelling spans a wide range of objectives and scales. The longest-standing thread is discriminative pretraining: Nucleotide Transformer [Dalla-Torre et al., 2025], DNABERT-2 [Zhou et al., 2024], and HyenaDNA [Nguyen et al., 2023] use masked or next-token pretraining and are evaluated primarily on downstream tasks such as variant-effect prediction. They expose no generative interface. Evo [Nguyen et al., 2024] introduced large-scale autoregressive pretraining over bacterial genomes, and Evo 2 [Brixi et al., 2026] extends the recipe to a 40B-parameter model trained across all domains of life with a small fixed catalogue of species tags that can be prepended to the context. From a designer’s perspective, Evo 2 is still a sequence completer: a specification like “a high-copy *E. coli* vector with kanamycin resistance and an EGFP reporter” has no representation in its prompt vocabulary. Decoder-only models trained on engineered plasmid corpora have recently been used for full-vector generation, including fine-tuning with wet-lab validation [Cunningham et al., 2025] and reinforcement-learning post-training that improves constraint satisfaction [Thiel et al., 2026]. Conditional generators that do offer some prompt interface have historically been restricted to short regulatory elements: 5^*′*^-UTR design from massively parallel expression assays [Sample et al., 2019], ribosome binding-site optimisation [Salis et al., 2009], discrete-diffusion enhancer and promoter generation [Stark et al., 2024], and cell-type-conditional enhancer design [Chen et al., 2024, Taskiran et al., 2024]. These methods operate on isolated elements of ~ 100 to 1,000 bp and target either a single scalar objective (e.g. expression level, TF binding score) or a single contextual condition (e.g. cell type). The coordination constraints that arise when an entire multi-kilobase construct must simultaneously contain a compatible origin, selection marker, regulatory cassette, and payload are not addressed at this scale. In the protein domain, conditional generation is further along, with family-conditional models [Madani et al., 2023] and discrete-diffusion approaches [Alamdari et al., 2023] demonstrating controllable single-chain generation; the analogous problem for DNA at the construct level has remained open. We discuss the relationship to Evo 2 in Appendix H.

### 2.2 Instruction tuning and RL for sequence design

Outside biology, the recipe of post-training a pretrained autoregressive model to follow user specifications is now standard. Supervised instruction tuning on prompt-response pairs [Wei et al., 2022] produces models that generalise to unseen tasks, and reinforcement learning from human or automated feedback [Christiano et al., 2017, Stiennon et al., 2020, Ouyang et al., 2022] further aligns model behaviour with a desired specification. PPO-based RLHF, direct preference optimisation [Rafailov et al., 2023], and group-relative policy optimisation (GRPO) [Shao et al., 2024] are the dominant algorithmic variants; the latter has driven substantial gains on tasks where outcomes can be *verified* automatically, most prominently mathematical reasoning [Shao et al., 2024, Lightman et al., 2024] and open-ended post-training recipes [Lambert et al., 2024]. The same paradigm has begun to migrate into biological sequence design: GFlowNets for protein generation [Jain et al., 2022], RL for small-molecule drug design [Zhou et al., 2019], foundation-model-based RNA inverse folding [Yang et al., 2024], and RL-driven optimisation of cis-regulatory elements [Taskiran et al., 2024]. The reward signals there span scalar property models, folding-energy and structure-prediction surrogates, and oracle assays; the unifying feature is differentiable or verifiable scoring on a single short sequence. Our contribution sits in parallel to text language modelling rather than to those sequence-design lines: we apply the same supervised-then-reinforced post-training pattern that turns a text LM into an instruction follower to a DNA LM, where the specification is a list of named components, the reward is a sequence-annotation pipeline, and the unit of generation is a multi-kilobase construct rather than a short element.

## 3 Methods

### 3.1 Dataset construction

Plasmid sequences were drawn from the Addgene repository [Addgene, 2024], yielding a corpus of 108,468 unique prompt–sequence pairs after deduplication and length filtering. These pairs span 20,644 unique plasmid IDs and 5,124 unique prompts; each plasmid contributes multiple prompt–sequence pairs reflecting different valid annotation subsets. We split 95%/5% into 103,045 training and 5,423 validation pairs (by random split, seed 42). A separate holdout of 22,400 prompt– sequence pairs, disjoint from pretraining, provides the pool from which the 1,000-prompt evaluation benchmark is sampled. Each sequence was annotated using pLannotate [McGuffie and Barrick, 2021], which queries four curated databases: the GenoLIB/SnapGene feature database (BLASTN), FPbase fluorescent proteins and Swiss-Prot proteins (DIAMOND), and Rfam non-coding RNA families (Infernal covariance models).

### 3.2 Special token design

Lab scientists describe plasmids at two granularities: named parts where exact identity matters operationally (ampicillin and chloramphenicol resistance are not interchangeable in a clone), and directional categories where designers think in classes rather than parts (high vs. low copy, mammalian vs. bacterial). The vocabulary mirrors this split, and the split aligns with what an alignment-based oracle can verify. *Hard tokens* (57 types in seven categories: `ORI_, AMR_, PROM_, ELEM_, REPORTER_, REP_, TAG_;` e.g. `<ORI_COLE1>, <AMR_AMPICILLIN>, <REPORTER_GFP>)` name sequence-verifiable components and are the target of both the GRPO reward and the faithfulness metric. *Soft tokens* encode metadata that no alignment can confirm — vector type (`VEC_`), host species (`SP_`), copy-number class (`<COPY_HIGH>, <COPY_LOW>`), and backbone identity (`BB_`) — and appear in training prompts but are excluded from the reward and the fidelity score.

A *motif registry* bridges human-readable hard-token names to pLannotate’s gene-level identifiers (e.g. `AMR_AMPICILLIN AmpR, → AmpR_(2)`, …), mapping each of the 57 token types to all qualifying sequence variants (codon-optimised forms, truncated cloning artefacts, source-organism variants). After filtering at≥ 95% identity and ≥80% coverage, the registry contains 660 entries, with each token type attested in at least 309 training pairs. The same registry defines the GRPO reward and the evaluation fidelity metric, ensuring evaluation directly measures what was reinforced. Each training example takes the form

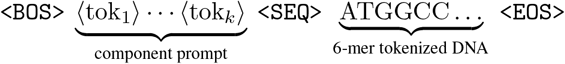

with component tokens sorted canonically; `<SEQ>` marks the prompt-to-sequence transition.

### 3.3 Model architecture and pretraining

PlasmidLM is a decoder-only transformer with 19.3 million parameters: hidden dimension 384, 10 layers, 8 attention heads of head dimension 48, rotary positional embeddings (RoPE; Su et al., 2024), RMSNorm [Zhang and Sennrich, 2019], and weight tying [Press and Wolf, 2017] between the input embedding and the language modeling head. A custom 6-mer tokenizer with stride 3 produces a vocabulary of 4,096 DNA k-mers plus special tokens, and a maximum positional context of 16,384 tokens supports the multi-kilobase plasmids in our corpus. Pretraining used standard next-token prediction on the training split (103,045 pairs) to step 65,000 (batch size 64, AdamW, cosine schedule peaking at 3× 10^*−*4^), with the released checkpoint reaching eval loss 0.129 and token accuracy 97.4%.

### 3.4 GRPO post-training

To sharpen prompt fidelity beyond what supervised pairing achieves, we apply Group Relative Policy Optimization (GRPO) [Shao et al., 2024]. For each training prompt, the policy generates *G* = 8 candidate completions. Each completion is scored by a composite reward: for every requested component token, BLAST is run against the corresponding motif registry reference sequences, and partial credit is awarded based on percent identity, alignment coverage, and normalized bit score. An aggregate recall penalty (Appendix A.1) ensures sequences that miss components entirely receive low reward regardless of per-hit quality. Rewards are group-normalized to compute advantages, and a clamped KL penalty (*β* = 0.5) applied to the frozen reference policy stabilizes training. Full hyperparameters are in Appendix A.

## 4 Evaluation framework

### 4.1 Two evaluation axes

#### Viable

A generated sequence is *viable* iff pLannotate [McGuffie and Barrick, 2021] detects an origin of replication, a selection marker, and at least one of a promoter or a coding sequence. Terminators and CDSs are recorded for descriptive purposes but do not gate viability. Let *V*_*i*_ ∈{0, 1} denote the viability indicator for prompt *i*’s generated sequence.

#### Faithful

For each hard token *t* requested in prompt *i*, the token is *recovered* iff the set of pLannotate sseqids attributed to *t* by the motif registry intersects the set of sseqids that pLannotate detects in the candidate sequence. The ≥95% identity and ≥80% coverage thresholds appear at *registry construction* time (Section 3.2), where they decide which reference variants qualify as canonical exemplars of *t*; per-prompt scoring is then a set-intersection on already-qualified sseqids, which is the same rule used by the GRPO reward. Per-prompt faithfulness is *F*_*i*_ = *n*_found,*i*_*/n*_requested,*i*_, defined for prompts with *n*_requested,*i*_ *>* 0 (996 of 1,000).

#### Useful-plasmid rate

The conjunction of the two axes is the headline summary metric. A sequence is *useful at threshold T* iff it is viable *and* achieves 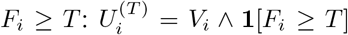. Viability alone (equivalently *T* =0) cannot distinguish a sequence that ignores the prompt from one that follows it; faithfulness alone rewards sequences that concatenate the requested motifs on an unstable backbone. Throughout we use *T* =0.5 as the default summary threshold because at-least-half of requested components present is a reasonable user-facing minimum for a prompt-following system; the threshold sweep in Appendix B reports the full range and the GRPO advantage at every positive *T*. The GRPO advantage is significant at every practical threshold (Table 4); *T* =0.5 is a representative operating point rather than a tuned one.

### 4.2 Annotation pipeline and motif registry

Sequences are annotated with pLannotate (named-feature detection via BLASTN, DIAMOND, and Infernal against four curated databases [McGuffie and Barrick, 2021]), Prodigal [Hyatt et al., 2010] (ORF prediction in metagenomic mode), and dustmasker [Morgulis et al., 2006] (low-complexity intervals). The motif registry operates at two levels: 660 sequence-level entries (individual reference variants) grouped into 57 component types (named parts such as ORI_COLE1 or AMR_AMPICILLIN); see Section 3.2. Running this exact pipeline on real training plasmids recovers 71.8% of requested tokens (token-pooled across prompts), establishing a practical detection ceiling that bounds any model’s faithfulness score from above. The remaining 28.2% are tokens whose references are too short to clear BLAST’s bit-score threshold (most epitope tags), or whose codon-optimised variants fall below the identity threshold. The viability rate on the same real-plasmid panel is 95.0% under our viability rule. Faithfulness numbers in this paper are therefore most usefully read against these two same-pipeline ceilings rather than against an idealised 100%.

### 4.3 Sampling protocols

We evaluate generations under two sampling regimes. The *single-shot* regime draws one sample per prompt at temperature 0.7, top-*p*=0.95, top-*k*=50, and is the default for headline numbers. The *best-of-N* regime draws *N* independent samples per prompt and keeps the candidate with the highest annotation-pipeline *n*_found_ score, breaking ties by viability then faithfulness. This is realistic for users with access to the same annotation pipeline at inference time, who can run pLannotate on candidates and reject failures; we report *N* ∈{1, 2, 4}. Where we contrast the post-trained model with the supervised base, both checkpoints generate one sample per prompt on the same 1,000 held-out prompts under identical sampling hyperparameters, supporting paired statistical tests on prompt-level outcomes. Note that the GRPO rollout temperature during post-training was 0.3 (Table 3), while evaluation uses temperature 0.7; the base-model temperature sweep in Appendix B.2 confirms that this evaluation setting is near the base model’s fidelity optimum.

## 5 Results

### 5.1 The useful-plasmid rate

PlasmidLM produces useful plasmids in 48.5% of single-shot generations and in 89.7% of best-of-4 generations on the held-out 1,000-prompt benchmark (Table 1, Figure 4). Random and dinucleotide-shuffled controls score 0.0%. The same annotation pipeline finds that 95.0% of held-out real Addgene plasmids are structurally plausible; the post-GRPO plausibility curve passes that reference at best-of-4 (95.8%, Figure 4 left), so on the viability axis post-training closes the gap to natural plasmids when the user can afford four samples per prompt.

**Table 1:** Useful-plasmid rate (*T* =0.5) on the held-out 1,000-prompt benchmark. Single-shot: temperature 0.7, top-*p*=0.95, top-*k*=50. Best-of-*K*: oracle ranker over *K* draws, four seeds. Real Addgene plasmids were not generated from prompts, so a useful-plasmid rate is undefined for them; the pipeline reports 95.0% structural plausibility on the real panel and 0.0% on shuffled controls.

|  | Single-shot | Best-of-2 | Best-of-4 |
| --- | --- | --- | --- |
| PlasmidLM (post-GRPO) | **48.5%** | **73.6%** | **89.7%** |
| PlasmidLM (supervised base) | 38.7% | 60.4% | 78.9% |
| Random GC-matched / shuffled | 0.0% | — | — |

The supervised base, before any verifiable-reward post-training, is already a useful model: it produces useful plasmids 38.7% of the time at single-shot decoding and 78.9% at best-of-4. Post-training adds +9.8 pp at single-shot decoding (paired bootstrap 95% CI [+5.7, +13.9], McNemar *p*≈6×10^*−*6^) and preserves a +10.8 pp advantage under best-of-4 (*p*≈2 ×10^*−*12^). The gain is significant at every positive faithfulness threshold and peaks near *T* =0.5 (threshold sweep in Appendix B). The gain is also not explained by sampling temperature: a sweep of the base model shows faithfulness peaking at *t*=0.3 and degrading at higher temperatures, so the *t*=0.7 evaluation setting already puts the base past its fidelity optimum; the GRPO advantage is not an artefact of a temperature that disadvantages the base (Appendix B.2).

### 5.2 Distributional and biophysical structure

The viable/faithful gate metrics treat each plasmid as pass-or-fail. We complement them with distributional checks against real Addgene at the population level, on properties the gate metric does not directly examine: length, GC content, codon usage in predicted ORFs, GC skew variation, ViennaRNA [Lorenz et al., 2011] minimum-free-energy density, and pairwise 6-mer diversity (Figure 3).

**Figure 3:**
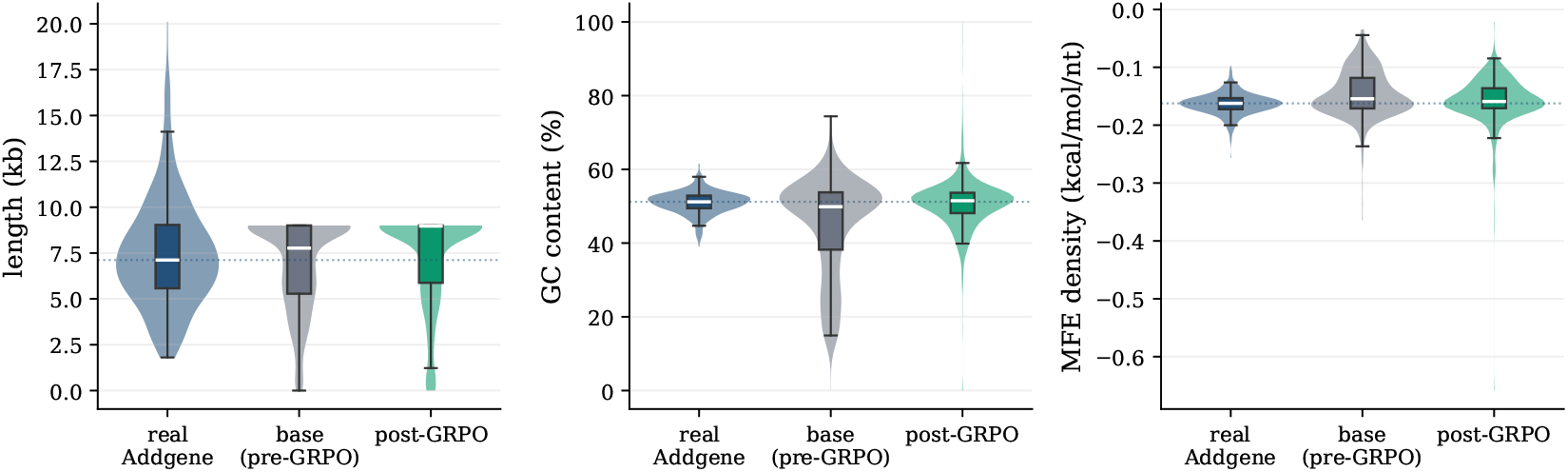
Length (kb), GC content (%), and ViennaRNA MFE density for real Addgene (*n*=500, random held-out subset), supervised base (*n*=1,000), and post-GRPO (*n*=1,000). IQR box with median; dotted line is real-population median. The GC panel shows the headline failure: a near-all-AT tail in the base that post-training truncates.

**Figure 4:**
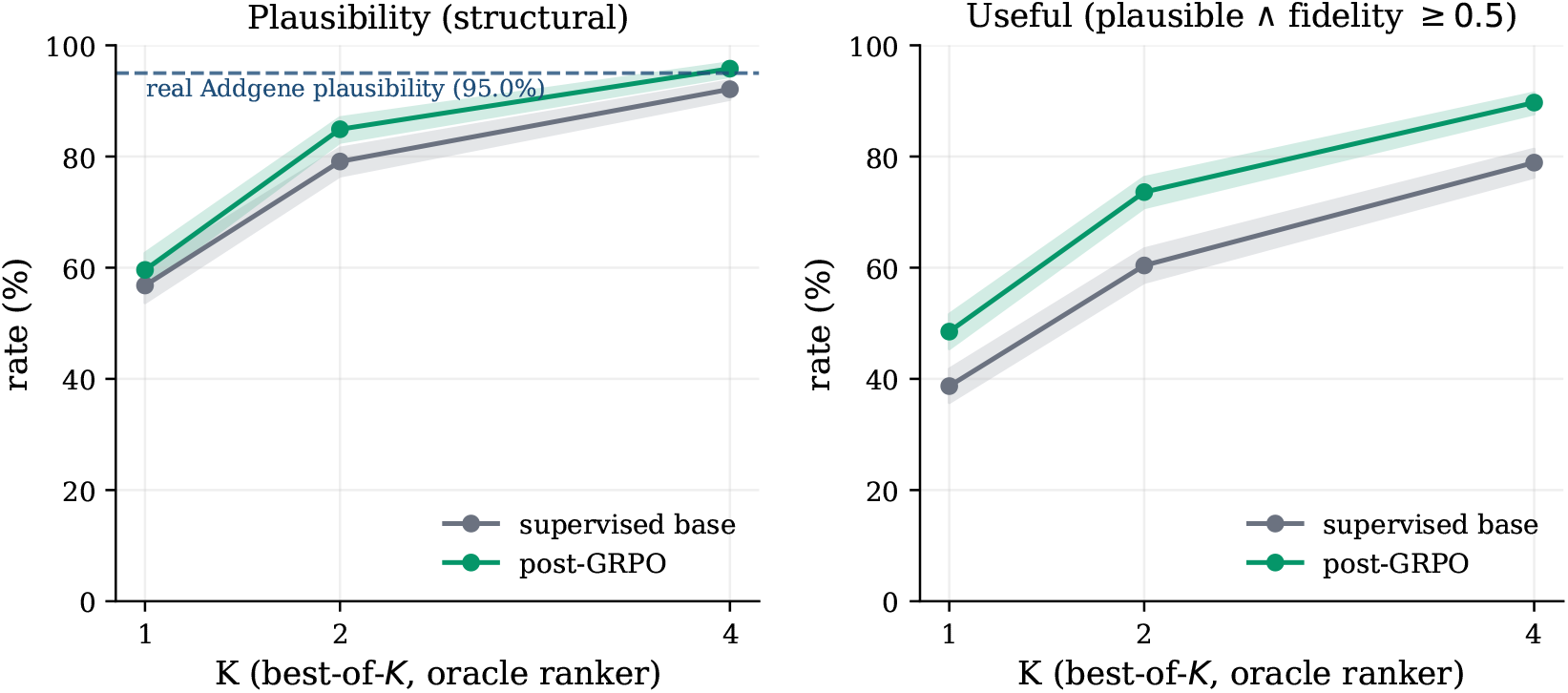
Best-of-*K* (*K*∈{1, 2, 4}), oracle ranker, *n*=1,000. Left: structural plausibility (dashed = real-Addgene 95.0%). Right: useful-plasmid rate (*T* =0.5). Wilson 95% bands shaded. Post-GRPO is strictly above the base at every *K*; plausibility reaches 95.8% at *K*=4. Useful-rate gap: +9.8 to +13.2 pp across *K*.

Mean lengths are 6.9 kb (base) and 7.2 kb (post-GRPO), against an Addgene reference of 7.5 kb. The more informative pattern is in GC content: the supervised base produces a long low-GC tail with mean 45.0% driven by a subpopulation of near-all-AT outputs, absent from real plasmids; post-training truncates this tail and recovers a real-like distribution with mean 50.6% (real Addgene mean 51.0%). MFE density [Lorenz et al., 2011] tells the same story: real plasmids sit at − 0.163 kcal/mol/nt, the supervised base at−0.146, and post-GRPO at −0.158 — a 3.1% relative gap from the real value (|−0.158 −(− 0.163) |/| −0.163|) (per-sequence ViennaRNA fold under DNA Mathews 2004 parameters; sequences over 5 kb are summarised as the mean MFE density across 5 random 2 kb windows; see Appendix I). Codon-usage JSD against *E. coli* is 0.131 in predicted open reading frames; GC skew variation is 0.0566 versus reference 0.0593. Generated sequences are at least as diverse as real plasmids on 6-mer statistics [Brown and Irber, 2016], with zero sourmash containment hits above threshold 0.3 in either run, providing no evidence of wholesale training-set memorisation. A LightGBM discriminator separating generated from real plasmids on sequence-level features achieves AUC 1.000 across all four base seeds, indicating that supervised-base outputs are perfectly distinguishable from real plasmids. Post-GRPO outputs are harder to separate (AUC 0.937 to 1.000 across four seeds), consistent with post-training recovering distributional structure that the base misses, though seed-level variance limits the strength of this conclusion. Post-training therefore recovers coarse distributional structure that the supervised base misses; the next subsection looks at finer-grained codon-level structure conditioned on the prompt.

### 5.3 Host-conditional codon usage

A registry-independent test of soft-token conditioning: for each generated and real plasmid we compute the codon adaptation index [Sharp and Li, 1987] against *E. coli, S. cerevisiae*, human, and mouse codon-usage tables [Nakamura et al., 2000], and summarise the cleanest contrast as ∆ = CAI_yeast_ −CAI_ecoli_ per sequence. Real plasmids show the expected diagonal across all four host classes. The post-GRPO model direction-matches real plasmids on three of four (`SP_ECOLI, SP_HUMAN, SP_MOUSE`) and shows a non-significant reversal on one (`SP_YEAST`, ∆ = −0.008 [−0.018, +0.002], where the real direction is positive). The supervised base direction-matches on two of four. A second pattern is that the supervised base produces sequences with higher matched-host CAI than typical real plasmids; post-GRPO normalises these back toward real-distribution values. Full methodology, sample sizes, and Figure 13 are in Appendix F.

### 5.4 Where prompt fidelity fails

Looking inside the faithfulness signal exposes both what the model can and cannot do. Per-category recovery rates (Table 2) show post-training improving five of six categories with gains ranging from +2 pp on reporters to +7.8 pp on origins of replication, and regressing on epitope tags from 8.4% to 2.8%. Epitope tags are short peptide motifs of 6 to 18 bp that BLAST rarely detects above the registry’s identity and coverage thresholds, so the training-time reward seldom credits them; the post-trained policy deprioritises them in proportion. We return to this in the discussion as an instance of reward misspecification under automated annotation.

**Table 2:**
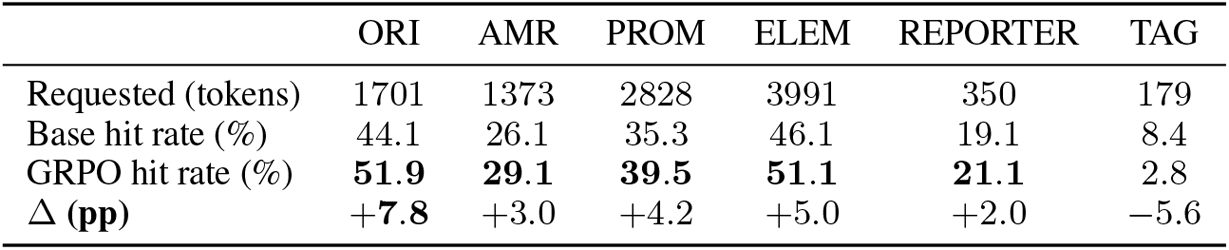
Per-category hard-token faithfulness on the 1,000-prompt evaluation. Cells are token-pooled hit rates (recovered/requested within category). REP tokens (*<* 1% of prompts) omitted. Per-prompt mean 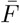in the text uses a different aggregation (mean of *F*_*i*_ over prompts) and will differ slightly.

|  | ORI | AMR | PROM | ELEM | REPORTER | TAG |
| --- | --- | --- | --- | --- | --- | --- |
| Requested (tokens) | 1701 | 1373 | 2828 | 3991 | 350 | 179 |
| Base hit rate (%) | 44.1 | 26.1 | 35.3 | 46.1 | 19.1 | 8.4 |
| GRPO hit rate (%) | **51.9** | **29.1** | **39.5** | **51.1** | **21.1** | 2.8 |
| $\Delta$ (pp) | **+7.8** | +3.0 | +4.2 | +5.0 | +2.0 | -5.6 |

The category-level view hides a strong long-tail effect at the token level (Appendix Figure 9). A handful of high-frequency tokens (`ELEM_IRES, ELEM_TRACRRNA, ORI_COLE1, PROM_AMPR`) achieve 50 to 60% hit rates, while a long tail of rarer tokens stays at or near zero (`AMR_CHLORAMPHENICOL 0/82, ORI_P15A 0/21, ORI_PSC101 0/17, ORI_RSF 0/8`, and most short epitope tags). The failure mode is a training-distribution artefact: tokens that appear fewer than~100 times in pretraining are effectively outside the model’s actionable vocabulary, regardless of post-training. The same long tail explains both the TAG regression and the `COPY_LOW` soft-token failure in Section 5.5. A barchart visualisation with Wilson 95% intervals appears in Appendix B (Figure 6).

**Figure 5:**
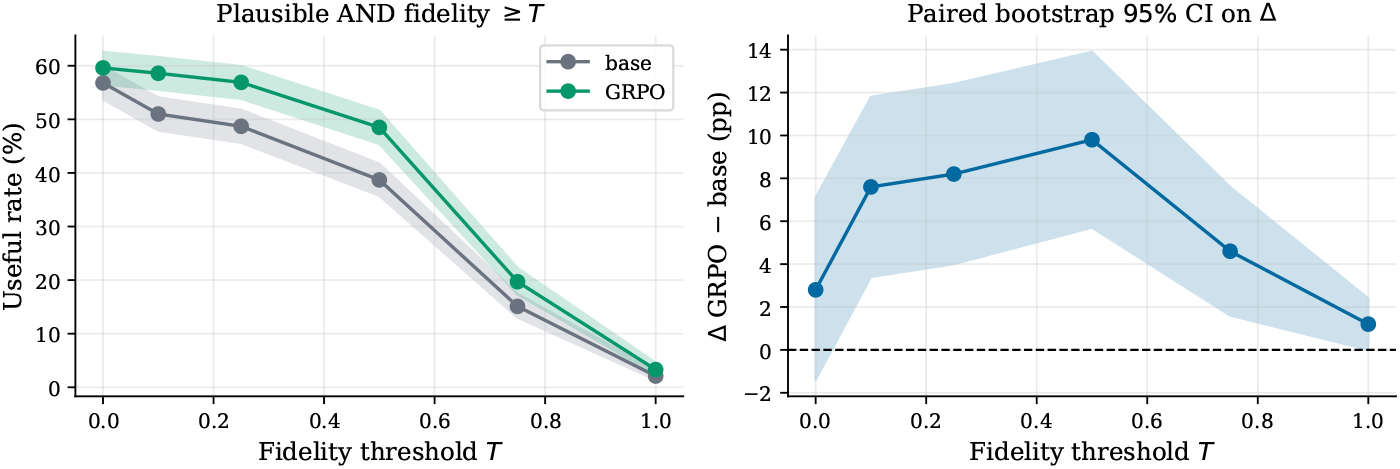
“Faithful plasmid” rate vs. fidelity threshold *T* for both checkpoints with paired bootstrap 95% bands (left), and the paired GRPO −base delta across *T* (right). The effect peaks near *T* = 0.5 and is significant at every positive threshold.

**Figure 6:**
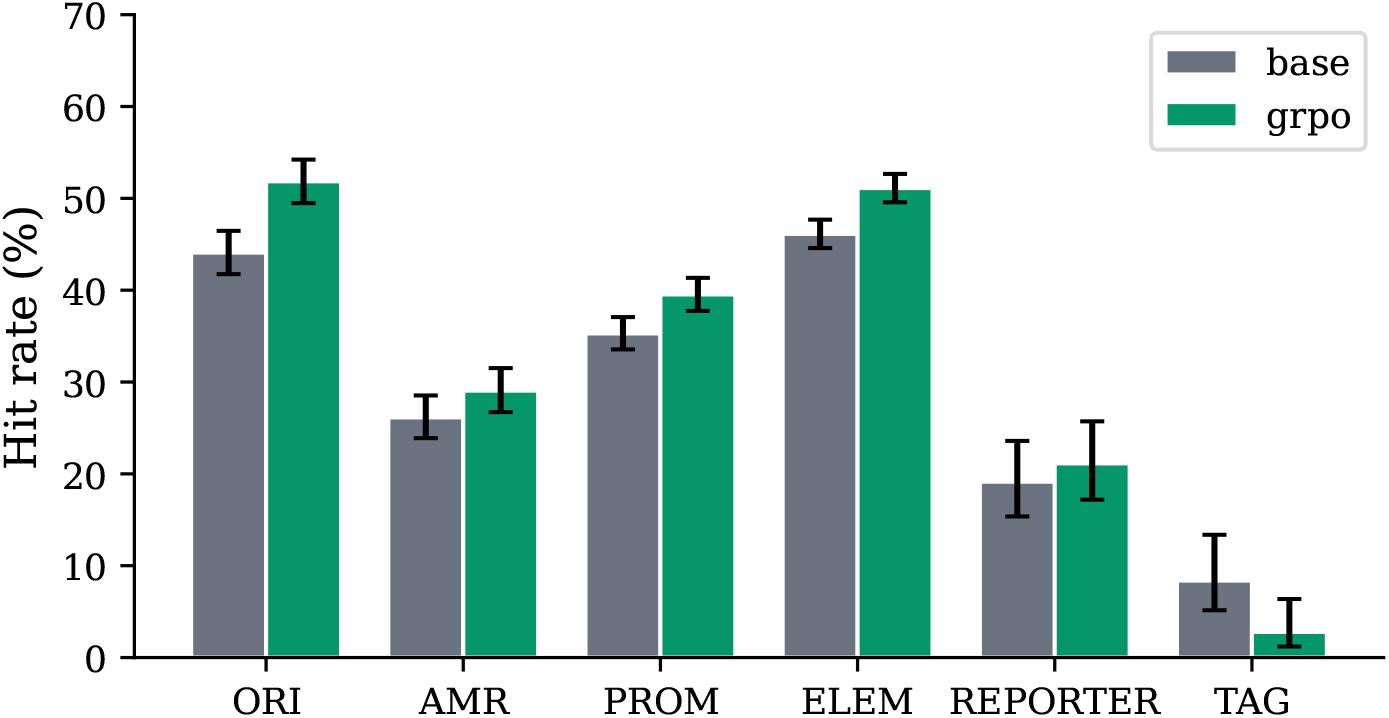
Per-category prompt faithfulness on the held-out 1,000-prompt benchmark, supervised base versus post-GRPO. Error bars are Wilson 95% intervals on pooled counts; see Appendix C for the non-independence caveat.

### 5.5 Soft-token conditioning

The prompt vocabulary includes *soft* metadata tokens, namely vector type (`VEC_*`), target species (`SP_*`), copy-number class (`COPY_{HIGH,LOW}`), and backbone identity (`BB_*`), that cannot be verified by sequence alignment and are excluded from the GRPO reward. Conditioning on soft tokens is therefore a test of pretraining alone, with post-training serving as a control that should not move the needle. Results split cleanly: tokens with biologically exclusive signatures are conditioned on reliably (VEC_LENTIVIRAL ∆ ≈+0.30, *p*≤ 10^*−*10^; `VEC_AAV` ∆ ≈+0.17, *p*≤ 6 ×10^*−*4^), while rare-class tags fail by the same long-tail mechanism as rare hard tokens: `COPY_LOW` is effectively ignored (the model returns a high-copy origin 42 to 63% of the time when a low-copy origin is requested), and post-training does not rescue any failure because soft tokens are absent from the reward. Full Fisher’s exact tables, forest plots, and COPY-class breakdowns are in Appendix E.

## 6 Discussion

The main result is that a small, specialist DNA language model can be promptable. A designer hands the model a list of named components in human-readable form and receives a multi-kilobase candidate construct that is viable and faithful most of the time at single-shot decoding, and almost all of the time under cheap rejection sampling against the same automated annotation pipeline. The viable/faithful separation is what made this measurable: viability alone hides whether the output is on-prompt, and per-token recovery alone hides whether the surrounding sequence is a plausible plasmid. Their conjunction, the useful-plasmid rate, is the metric we recommend as default for promptable construct design.

### 6.1 Failure modes of automated-feedback post-training

The most informative failure is the epitope tag regression (`TAG_*: 8.4%→2.8%, −5.6 pp`). TAG tokens are short peptide motifs of 6 to 18 bp that BLAST struggles to detect even in real plasmids; the reward signal on a correct TAG is weak and noisy. The policy responds as RL-from-automated-feedback analyses predict [Skalse et al., 2022, Krakovna et al., 2020]: it deprioritises what the scorer cannot credit. The zero-hit rare tokens (Appendix D; `AMR_CHLORAMPHENICOL` 0/82, `ORI_P15A` 0/21, `ORI_PSC101` 0/17) show the same pattern at vocabulary scale: rare-class tokens are pushed toward zero even when they appear in the prompt. This is the primary risk for any practitioner adopting this recipe, and it has a clear diagnostic: per-token hit-rate audits before and after post-training reveal exactly which parts of the prompt vocabulary the reward can and cannot see. The same effect appears at vocabulary scale in the zero-hit tail (Appendix D), affecting components that are common laboratory workhorses including chloramphenicol resistance, P15A origins, and low-copy backbones. Practitioners adopting the model should pre-screen the per-token hit-rate audit before deployment. Two mitigations are apparent: complementing BLAST scoring with HMM profiles or exact *k*-mer matching for short motifs, and rebalancing the training distribution to increase coverage of rare-class tokens.

### 6.2 Limitations

The model and its evaluation are both shaped by what an alignment-based annotation pipeline can see. The 71.8% faithfulness and 95.0% viability ceilings on real Addgene plasmids (Section 4.3) bound what any model can score; components without strong sequence-level signatures are difficult for the pipeline to credit and therefore difficult for the model to learn. Roughly a quarter of the hard-token vocabulary is effectively unreachable: tokens with fewer than ~100 training appearances stay near zero hit rate regardless of post-training (Section 5.4). Soft-token conditioning is partial rather than uniform: it works where biology supplies a distinctive sequence signature and fails where it does not (Section 5.5). The training corpus inherits Addgene’s bias toward common research vectors (low-copy origins, chloramphenicol resistance, and P15A backbones are underrepresented) and underrepresents therapy and industrial constructs.

#### Reward and metric share a registry

The GRPO reward and the faithfulness metric are both defined against the same 660-entry motif registry. The post-training delta is evidence that the model fits the scorer better; it is not by itself a model-independent claim of learned controllability.

All sequences are represented and generated as linear strings, although plasmids are biologically circular. The model has no explicit representation of circularity, so it cannot reason about features that span the linearisation boundary or enforce rotational equivalence between sequences that differ only in start position.

#### Biosafety and dual-use considerations

PlasmidLM generates sequences within the distribution of publicly deposited Addgene plasmids. The prompt vocabulary is bounded by the 57-token hard-token set derived from that corpus: there is no way to prompt the model toward novel harmful components, and no basis for assuming it can be sampled in a way that produces harmful outputs not already trivially accessible from the same public catalogue.

### 6.3 Future work

Three directions follow directly from the failure analysis. First, wet-lab validation: a small panel of generated constructs assembled and tested for whether the requested reporter expresses in the requested host. Second, a hybrid reward signal combining BLAST with HMM profiles and exact *k*-mer matching to recover the short motifs that current scoring misses. This is the direct fix to the TAG regression and the most actionable lesson from the per-token analysis. Third, closed-loop training in which the in-silico oracle is augmented or replaced by experimental feedback, with measured expression or assembly success serving as a reward signal. This is the natural extension once a wet-lab loop is established and is the path toward designer-aligned objectives that the current registry-bounded reward cannot express. Additional directions including distribution rebalancing, compositional prompting, and multi-objective rewards are discussed in Appendix G.

## 7 Conclusion

PlasmidLM is a promptable DNA language model: a designer specifies a multi-kilobase construct in human-readable terms and receives a candidate sequence in a single autoregressive pass. We propose the viable/faithful decomposition and the useful-plasmid rate as a primary evaluation framework for promptable DNA generation, and we report a useful-plasmid rate of 89.7% under best-of-4 sampling against the same annotation pipeline used for evaluation, and 48.5% at single-shot decoding, on a held-out 1,000-prompt benchmark. Verifiable-reward post-training with GRPO improves the useful-plasmid rate across all sampling budgets and fails in the way reward misspecification under automated annotation predicts. We release the model, evaluation suite, and paired benchmark to support replication and extension.

## A Model and training details

**Table 3:** Model and training hyperparameters.

| Parameter | Value |
| --- | --- |
| Hidden dimension | 384 |
| Layers | 10 |
| Attention heads | 8 |
| Head dimension | 48 |
| Total parameters | 19.3M |
| Vocabulary size | 4,096 + special tokens |
| Max positional context | 16,384 tokens |
| Positional encoding | RoPE |
| Normalization | RMSNorm |
| Weight tying | Yes (embed ↔ lm_head) |
| *Pretraining* |  |
| Optimizer | AdamW ($\beta_1=0.9$, $\beta_2=0.999$) |
| Peak learning rate | $3 \times 10^{-4}$ |
| Schedule | Cosine decay |
| Batch size | 64 |
| Steps (released checkpoint) | 65,000 |
| Eval loss | 0.129 (token accuracy 97.4%) |
| Hardware | 1 × A100-40GB |
| *GRPO post-training* |  |
| Group size ($G$) | 8 |
| Prompt batch size | 8 |
| Micro-batch size | 8 |
| Learning rate | $5 \times 10^{-6}$ |
| Warmup steps | 50 |
| KL coefficient ($\beta$, $k_3$ clamped) | 0.5 |
| Clip range | 0.2 |
| Scheduled steps | 1,000 |
| Released checkpoint | step 800 |
| Rollout temperature | 0.3 |
| Rollout top-$p$ | 0.95 |
| Rollout max new tokens | 2,500 |
| Reward | pLannotate composite (motif registry, sequence identity) |
| Hardware | 1 × GPU (L4/L40S/A100 pool, Anyscale Ray) |
| Training time | ∼17 hours |

**Table 4:** Composite “useful plasmid” rate (structurally plausible *and* fidelity ≥*T*) across thresholds on the paired *n* = 1000 evaluation. The GRPO gain is significant at every positive threshold; the effect peaks near *T* = 0.5.

| Threshold $T$ | Base | GRPO | $\Delta$ | paired 95% CI | McNemar $p$ |
| --- | --- | --- | --- | --- | --- |
| 0.00 (plausibility only) | 56.8% | 59.6% | +2.8 pp | $[-1.4, +7.1]$ | 0.21 |
| 0.25 | 48.7% | 56.9% | +8.2 pp | $[+4.0, +12.4]$ | $1.7 \times 10^{-4}$ |
| **0.50** | **38.7%** | **48.5%** | **+9.8 pp** | **$[+5.7, +13.9]$** | **$6.4 \times 10^{-6}$** |
| 0.75 | 15.1% | 19.7% | +4.6 pp | $[+1.7, +7.6]$ | $3.3 \times 10^{-3}$ |
| 1.00 (all tokens) | 2.1% | 3.3% | +1.2 pp | $[0.0, +2.4]$ | 0.065 |

### A.1 GRPO reward function

## B Additional results

### B.1 Useful plasmid rate across fidelity thresholds

### B.2 Base-model temperature sweep

## C Statistical conventions

Wilson 95% CIs on rates; paired bootstrap CIs (10,000 resamples, resampling prompts) on binary-outcome deltas; McNemar with continuity correction (exact when any cell *<* 25) on paired binary contingency tables. Per-sequence faithfulness CIs are bootstrapped over prompts because tokens within a prompt are not independent. The per-category hit-rate table reports raw token-pooled rates without CIs for the same reason; per-category Wilson intervals on pooled counts (used in Figure 6) are mildly optimistic and flagged as such. All artefacts are archived for public release.

## D Per-token hit-rate decomposition

## E Soft-token conditioning analysis

### Probe construction

For each soft value (e.g., VEC_LENTIVIRAL) we list the hard-token signature biology expects: LTRs, Ψ, cPPT, and WPRE in this case. Each hard token resolves to a set of pLannotate sseqids via the motif registry; the soft-value probe is the union of those sseqid sets. We then compare the probe-hit rate between prompts carrying the soft value and prompts carrying a different value in the same soft category, using Fisher’s exact (two-sided). Probes for tags without biologically exclusive signatures (e.g. VEC_MAMMALIAN, whose features appear in most plasmids for unrelated reasons) are included for completeness but are expected to wash out. Four soft categories are defined in the vocabulary: `BB_*` (20 values`COPY_{HIGH,LOW}, SP_* (9 values), VEC_*` (11 values).

## F Host-conditional codon usage (full analysis)

For each generated and real plasmid we predict open reading frames with Prodigal [Hyatt et al., 2010] and compute the codon adaptation index [Sharp and Li, 1987] of each predicted CDS against the codon-usage tables of *E. coli, S. cerevisiae*, human, and mouse (Kazusa [Nakamura et al., 2000], whole-CDS frequencies; CAI evaluated over body codons of ORFs≥ 100 codons, then length-weighted within each plasmid). The `SP_*` token in the prompt is a soft token: it is part of the input but never appears in the GRPO reward, so any host-conditional codon signal must come from supervised pretraining alone. A model that has learned host-specific codon usage should produce sequences whose CAI to a given host table tracks the host declared in the prompt. Sharp–Li CAI is GC-dependent and the four organisms have different GC content, so we summarise the cleanest contrast as ∆ = CAI_yeast_ −CAI_ecoli_ per sequence: yeast and *E. coli* have similar GC and different codon preferences, so this difference isolates the codon-preference signal from the GC confound.

**Figure 7:**
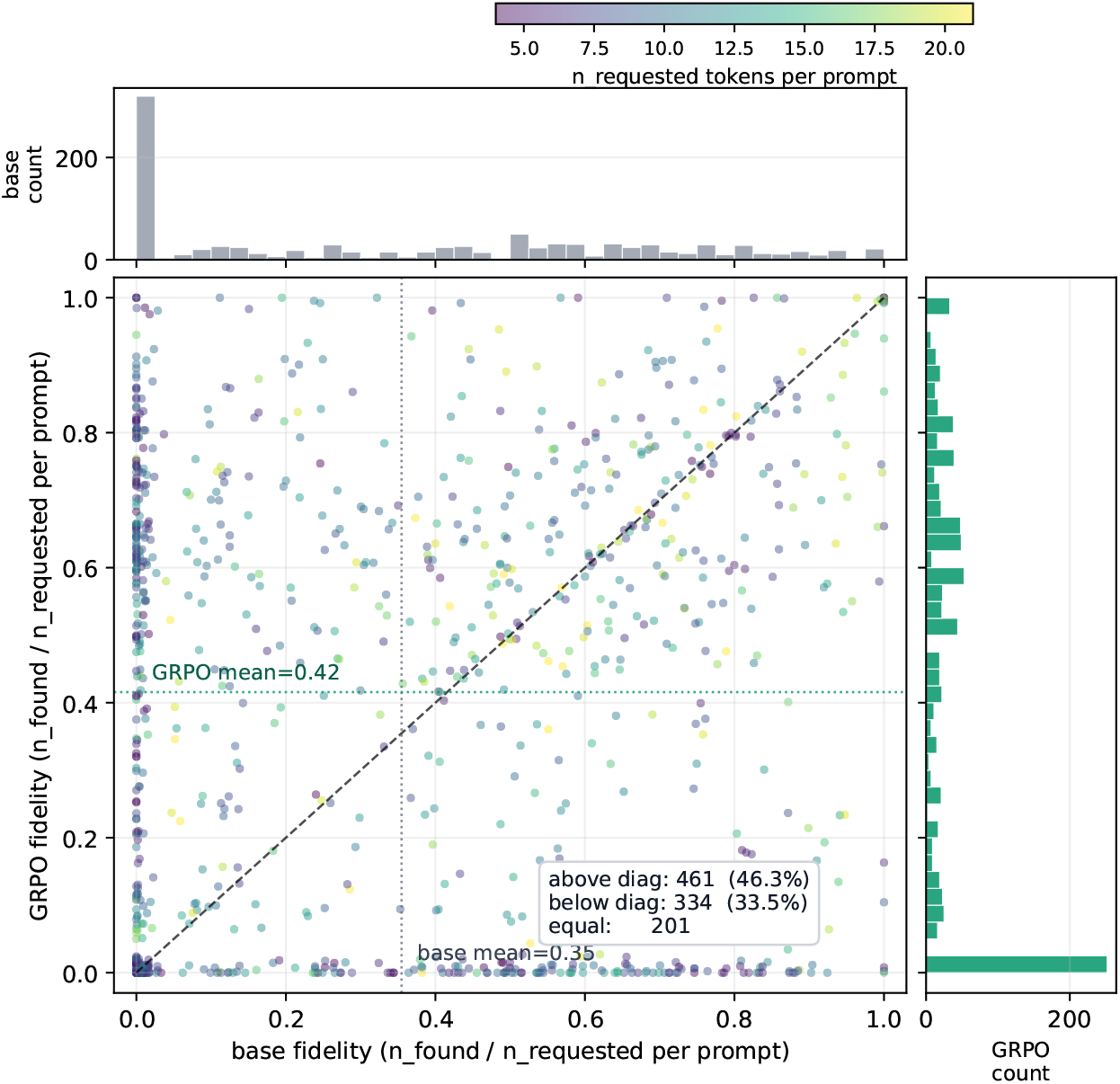
Per-prompt faithfulness on the held-out 1,000-prompt benchmark, supervised base (*x*-axis) vs. post-GRPO (*y*-axis). Marker jitter separates points landing on shared fractions (e.g. 1/2, 2/3). Point colour encodes the number of tokens requested per prompt; marginal histograms show each model’s univariate distribution. 46.3% of prompts improve under post-training, 33.5% regress, and 20.1% are unchanged; the improvement is broadly distributed across prompt complexity rather than concentrated on simple prompts.

Real plasmids show the expected diagonal across all four hosts (*n*=300 training plasmids per SP class, sampled with their original prompt strings): ∆ = −0.024 (95% CI [−0.035, −0.014]) for SP_ECOLI, +0.039 [+0.029, +0.049] for `SP_YEAST`, −0.006 [−0.012, − 0.001] for `SP_HUMAN`, and −0.013 [−0.019, −0.007] for `SP_MOUSE`. To get usable sample sizes for the rare `SP_ECOLI` and `SP_YEAST` classes (which appear in only a handful of prompts in the original 1,000-prompt evaluation), we generated *n*=400 additional sequences per checkpoint with prompts forced to those tokens, sampling with replacement from the 40 unique`SP_ECOLI` and 99 unique `SP_YEAST` training prompts. `SP_HUMAN` and `SP_MOUSE` use the existing eval-set generations, where these classes are well-represented.

Both checkpoints reproduce the diagonal partially (Figure 13). The post-GRPO model direction-matches real plasmids on three of four host classes (`SP_ECOLI, SP_HUMAN, SP_MOUSE`) and regresses on one (`SP_YEAST`, ∆ = −0.008 [−0.018, +0.002], where the real direction is positive). The supervised base direction-matches on two of four (`SP_YEAST` and `SP_HUMAN`), with the wrong sign on SP_MOUSE and a non-significant near-zero on SP_ECOLI. On the soft tokens GRPO never directly rewarded, post-training improves host-conditioning across the broader vocabulary even as it loses ground on yeast specifically. The `SP_YEAST` regression is the one note in the result we cannot account for and flag as a target for further study.

A second pattern, visible in absolute CAI rather than the contrast ∆, is consistent with the discriminator analysis. The supervised base produces sequences that read as more codon-optimised than typical real plasmids: matched-host CAI of 0.796 on `SP_HUMAN` prompts versus 0.732 for real Addgene of the same SP class, and similarly elevated values on `SP_ECOLI` (+0.022) and `SP_YEAST` (+0.054). Post-GRPO normalises these back: 0.745 on `SP_HUMAN` and 0.749 on `SP_MOUSE`, both within rounding of the real distribution; and on the bacterial and yeast classes, post-GRPO sits below real instead of above. The same direction the LightGBM discriminator sees (post-GRPO drops AUC from 1.000 towards 0.94) appears here at the codon level: the two observations are consistent with post-training reducing a generator-specific codon-optimisation signature inherited from pretraining, though we do not isolate this from other distributional shifts.

**Figure 8:**
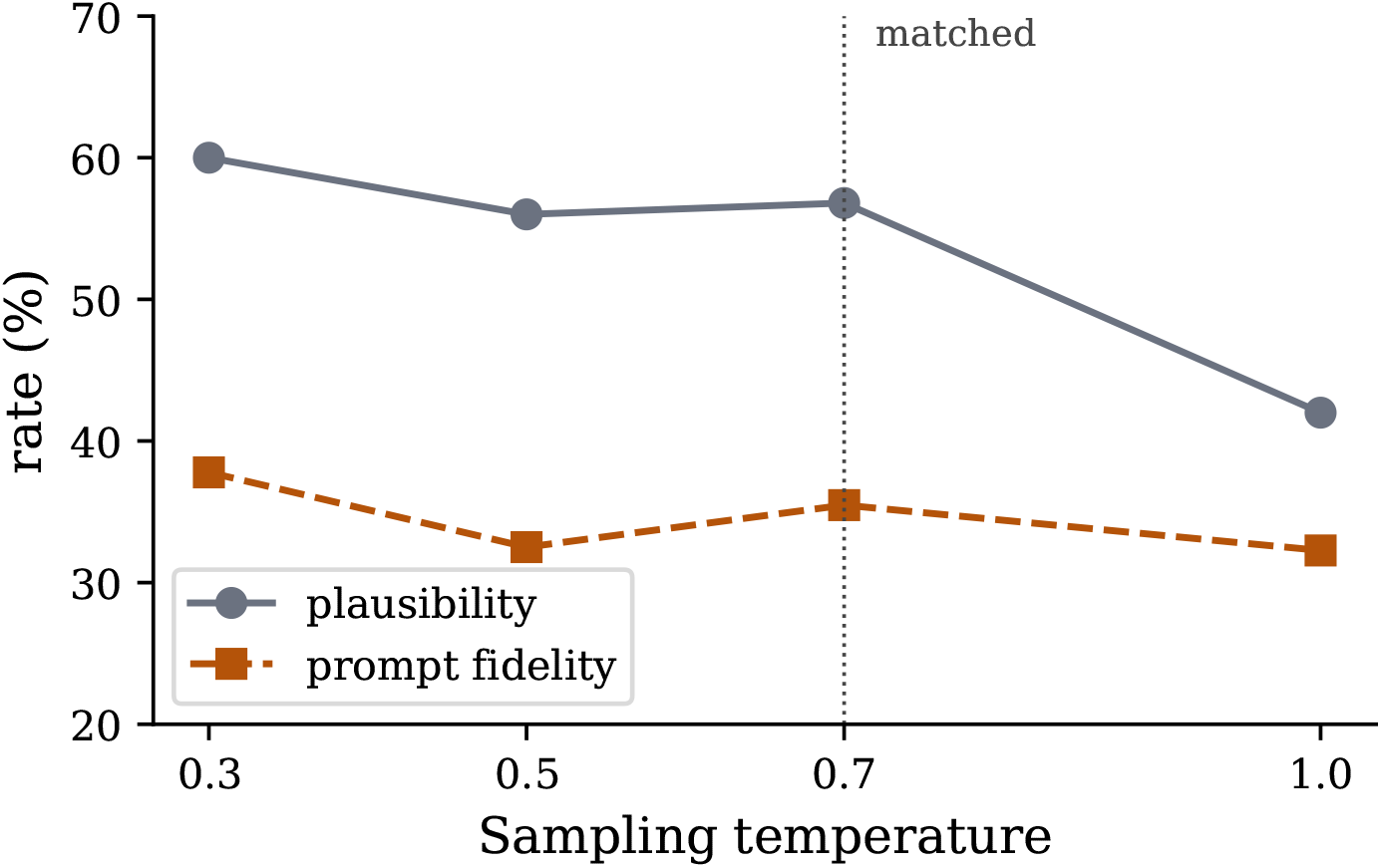
Pre-GRPO base model, small temperature sweep (*n* = 100 per cell; matched *n* = 1000 run for *t* = 0.7 for comparison). Base prompt fidelity peaks at *t* = 0.3 (37.8%) and degrades at higher temperatures; *t*=0.7 puts the base past its fidelity peak, so if anything the comparison is generous to the base. The GRPO advantage is not an artefact of an unfavourable temperature for the base.

## G Additional future directions

Distribution rebalancing or upsampling for rare-class tokens would lift the long-tail zero-hits. Compositional and counterfactual prompting (replace ampicillin with kanamycin in this construct; combine cassette A with backbone B) would test whether the model has learned compositional structure or memorised prompt-to-sequence mappings. The host-conditional codon-usage analysis revealed an unexplained `SP_YEAST` regression under post-training; a targeted ablation of training-set composition and reward-shape effects on this specific soft token is a concrete diagnostic. The recipe should generalise to AAV, lentiviral therapy, and CRISPR guide libraries, each of which has the three required ingredients (vocabulary, corpus, annotation pipeline). Multi-objective rewards combining motif faithfulness with predicted expression and dynamic range are a natural follow-up.

## H Discussion: broader context

### H.1 Relation to large generative DNA models

Evo 2 conditions on a small fixed catalogue of species tags but exposes no interface at the component level we study, so it cannot be prompted with the kinds of specifications PlasmidLM is designed to satisfy. A fair comparison would require either fitting a prompt interface onto Evo 2 or designing an evaluation framed around unconditional plausibility, both of which are out of scope here.

The choice of a small specialist over a fine-tuned generalist also has practical consequences worth stating explicitly. PlasmidLM trains in approximately 17 hours on a single GPU, runs inference cheaply, and fits on a workstation. The prompt vocabulary is purpose-built for the modality. Larger generalist DNA models offer no comparable component-level interface and no clear deployment story for the kinds of designer-facing workflows this work targets.

**Figure 9:**
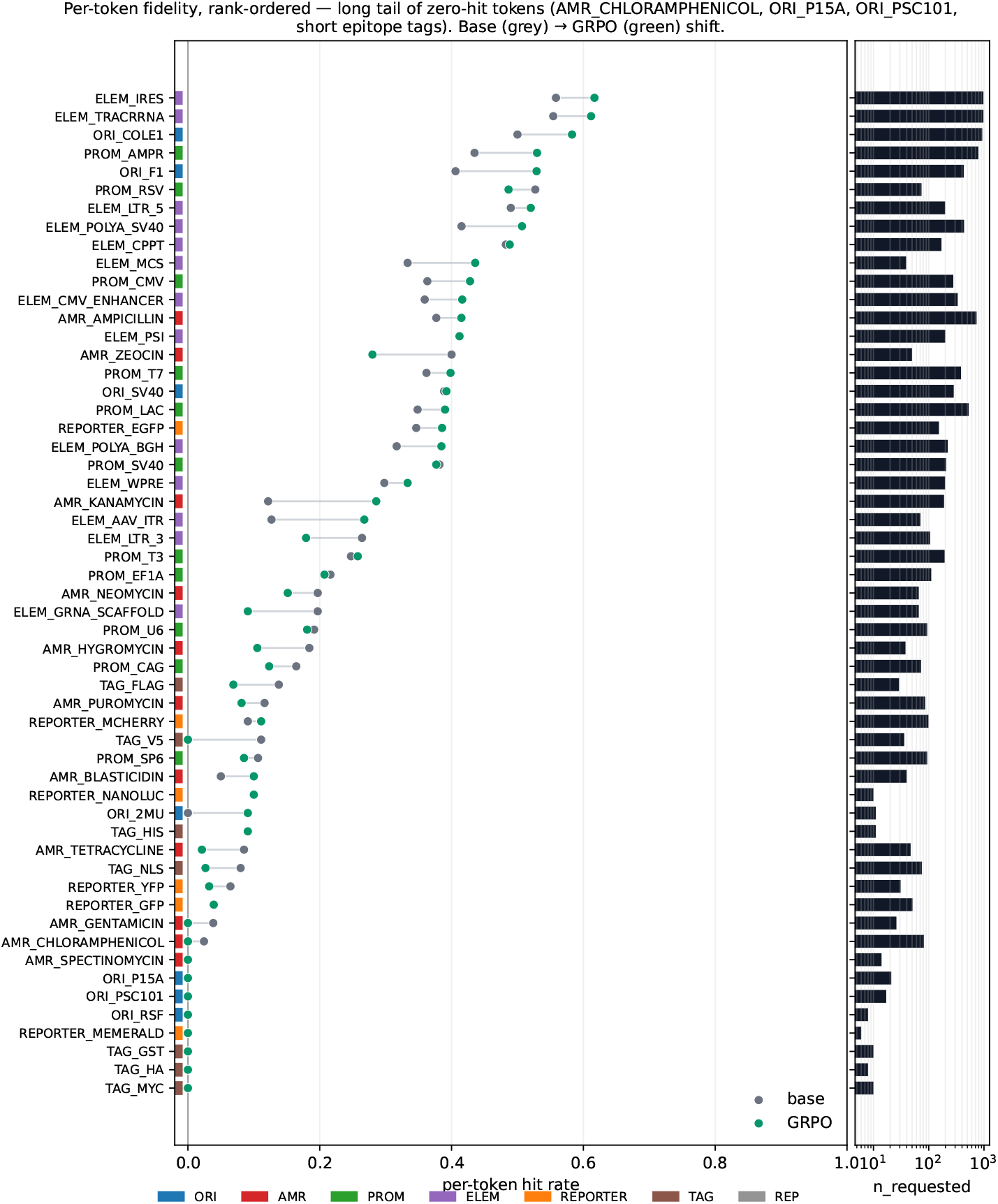
Per-token hit rate, rank-ordered by the maximum of the base and GRPO rates. Each row shows one hard token: base (grey) and GRPO (green) dots connected by a light line to highlight the shift. Category colour strip on the far left. Right panel is *n*_requested_ in the paired eval on a log scale. The figure makes two patterns visible: GRPO uniformly lifts the already-detectable tokens, and a long tail of rare tokens stays at zero in both runs (the zero-hit tail is the same set that explains the TAG regression and the `COPY_LOW` soft-token failure in Section 5.5).

### H.2 Beyond plasmids

The recipe used here, namely a curated component vocabulary, a small specialist language model, and verifiable post-training against a sequence-annotation pipeline, is not specific to plasmid biology. Any modality with (i) a designer-facing component vocabulary, (ii) a corpus of annotated examples, and (iii) an automated pipeline that detects those components in candidate sequences supports the same construction. Examples where each ingredient already exists include synthetic gene-circuit assemblies, AAV and lentiviral therapy vectors, and CRISPR guide-vector libraries, where the relevant components (promoters, regulatory cassettes, packaging signals, guide spacers) are detectable by existing alignment or HMM-based pipelines.

**Figure 10:**
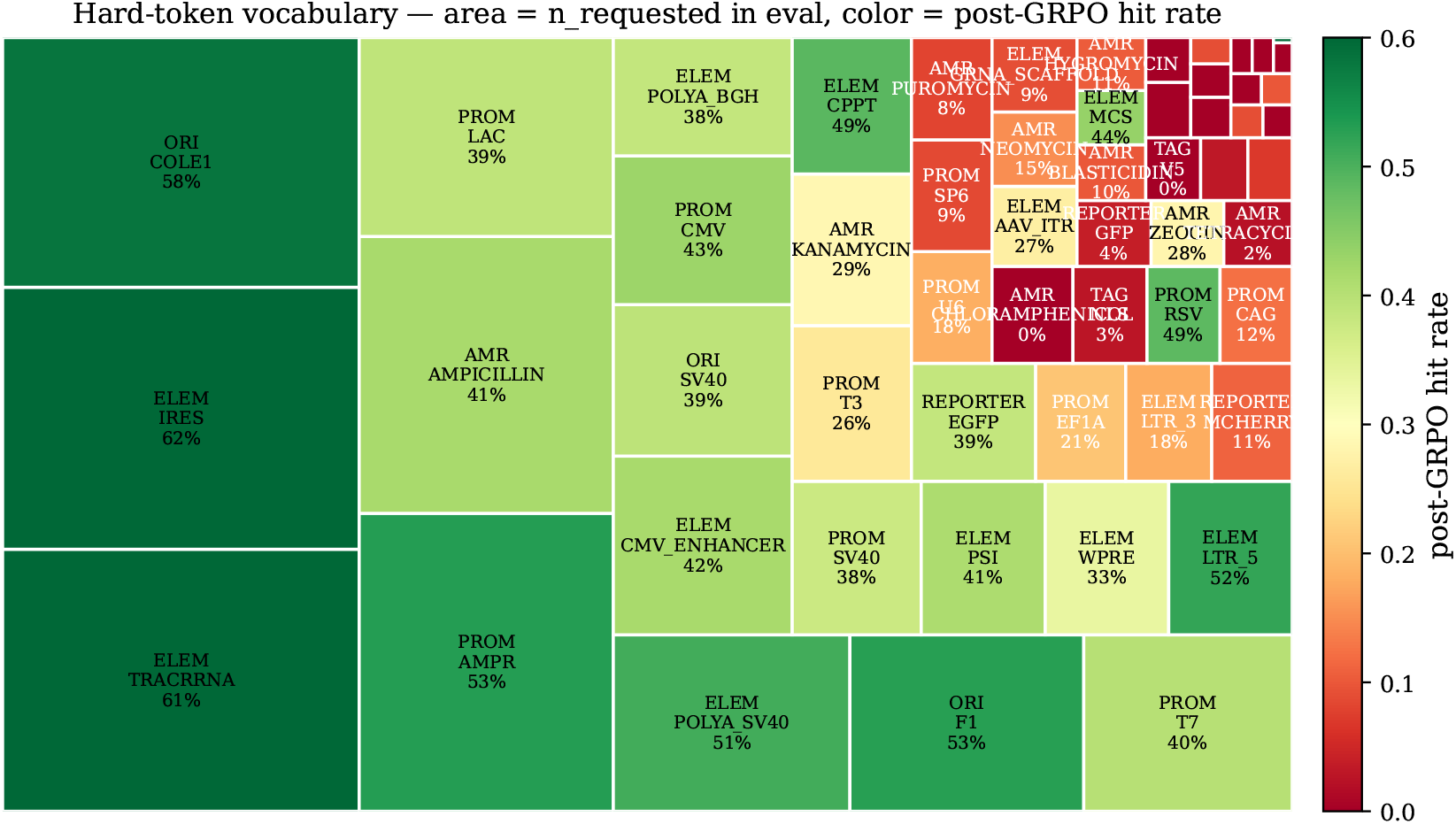
Hard-token vocabulary scope on the held-out benchmark, as a treemap. Tile area is the number of times the token appears in evaluation prompts; tile colour is the post-GRPO hit rate (0–60% red →green scale). The four large green tiles in the upper-left (`ELEM_TRACRRNA, ELEM_IRES, ORI_COLE1, PROM_AMPR`) carry most of the evaluation mass and achieve 50–58% hit rates. The long tail of small dark tiles on the right is the zero-hit set: rare antibiotic resistance markers, rare origins, and most short epitope tags. `REP_*` tokens have no evaluation prompts and are omitted. Companion to Figure 9, which shows base →GRPO movement; the treemap shows scope of vocabulary at a glance.

## I Evaluation pipeline and reproducibility

The complete evaluation pipeline constructs reference panels and negative controls, performs large-scale generation and annotation with pLannotate, Prodigal, and dustmasker, computes viability and fidelity metrics, trains the LightGBM discriminator, and computes best-of-*K* tables. All artifacts (generations, pLannotate JSON, per-prompt fidelity parquets, discriminator outputs) are archived for public release. Commands, random seeds, and environment specifications are included. The best-of-*K* analysis is runnable end-to-end as a PEP 723 script at `eval/scripts/analyze_best_of_k.py`; the paired marimo notebook `notebooks/base_vs_grpo_analysis.py` renders every figure and table in Section 5.1 directly from the archived data.

### Folding protocol

MFE density is computed with ViennaRNA’s fold under the DNA Mathews 2004 parameter set; sequences ≤5 kb are folded full-length, longer sequences are summarised as the mean MFE density across 5 random 2 kb windows.

### Model checkpoints

SHA-256 hashes for both checkpoints are recorded in the model card accompanying the release. Eval loss 0.129 and token accuracy 97.4% are recorded in the model card accompanying the base checkpoint; training logs were not retained.

**Figure 11:**
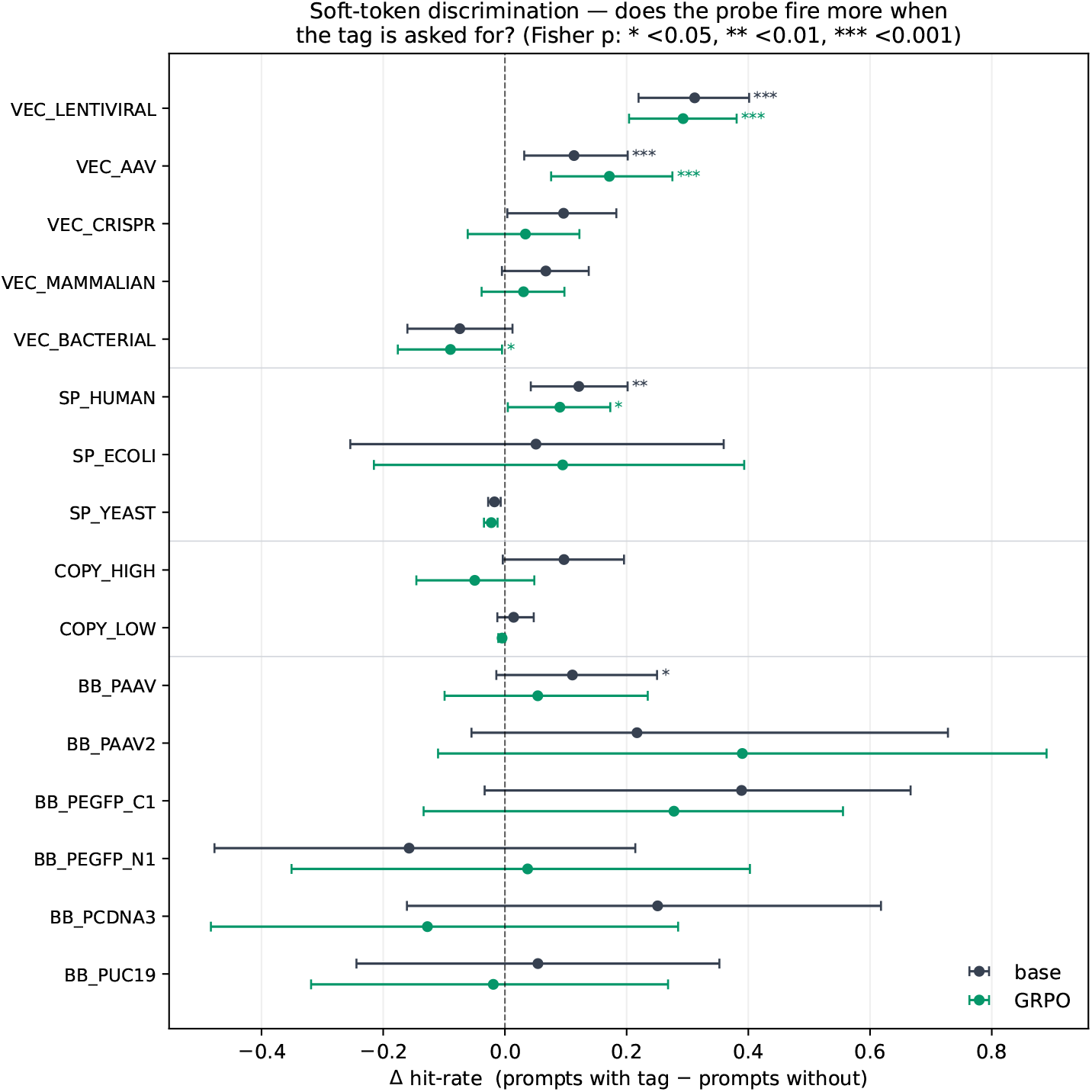
Soft-token discrimination. For each soft value with a biologically meaningful probe, ∆ is the difference in probe-hit rate between prompts that carry that tag and prompts that carry some other value in the same category; error bars are bootstrap 95% intervals; stars mark Fisher *p <* 0.05 (*), *<* 0.01 (**), *<* 0.001 (* * *). VEC_LENTIVIRAL and `VEC_AAV` show large, significant conditioning effects in both the pretrained base and the post-GRPO model. Most other tags are indistinguishable from zero; GRPO preserves but does not extend the pattern, consistent with soft tokens being absent from the reward.

**Figure 12:**
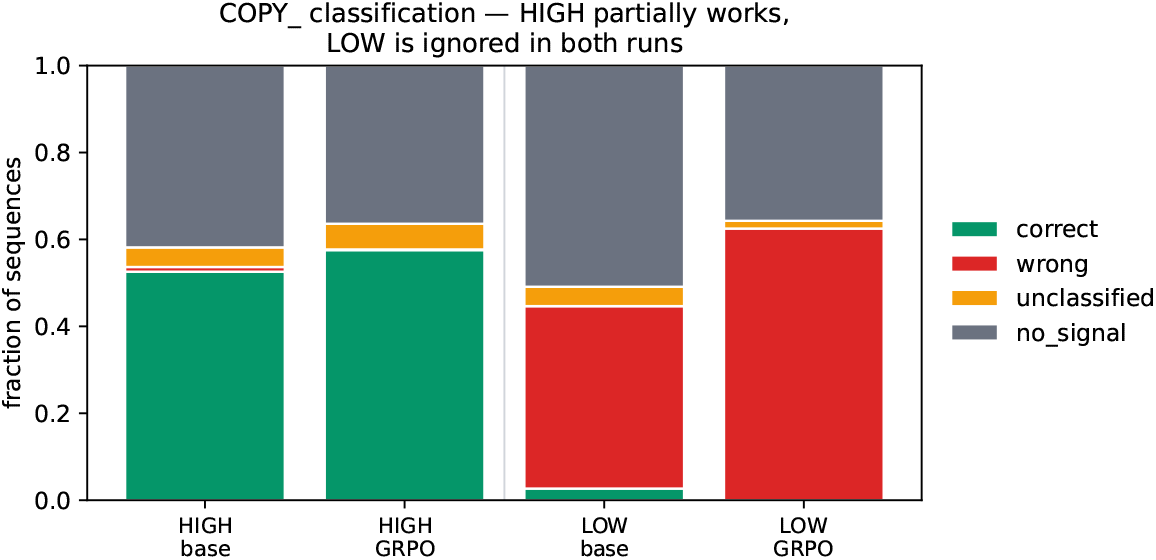
`COPY_` conditioning, stacked by outcome. *correct* = the requested copy class’s origin was detected; *wrong* = the opposing class’s origin was detected instead; *unclassified* = a non-discriminative origin (e.g. ORI_F1, ORI_SV40, ORI_2*µ*) was detected; *no_signal* = no origin detected. `COPY_HIGH` partially works in both runs; `COPY_LOW` is effectively ignored, with the model outputting a high-copy origin far more often than a low-copy one when low copy is requested.

**Figure 13:**
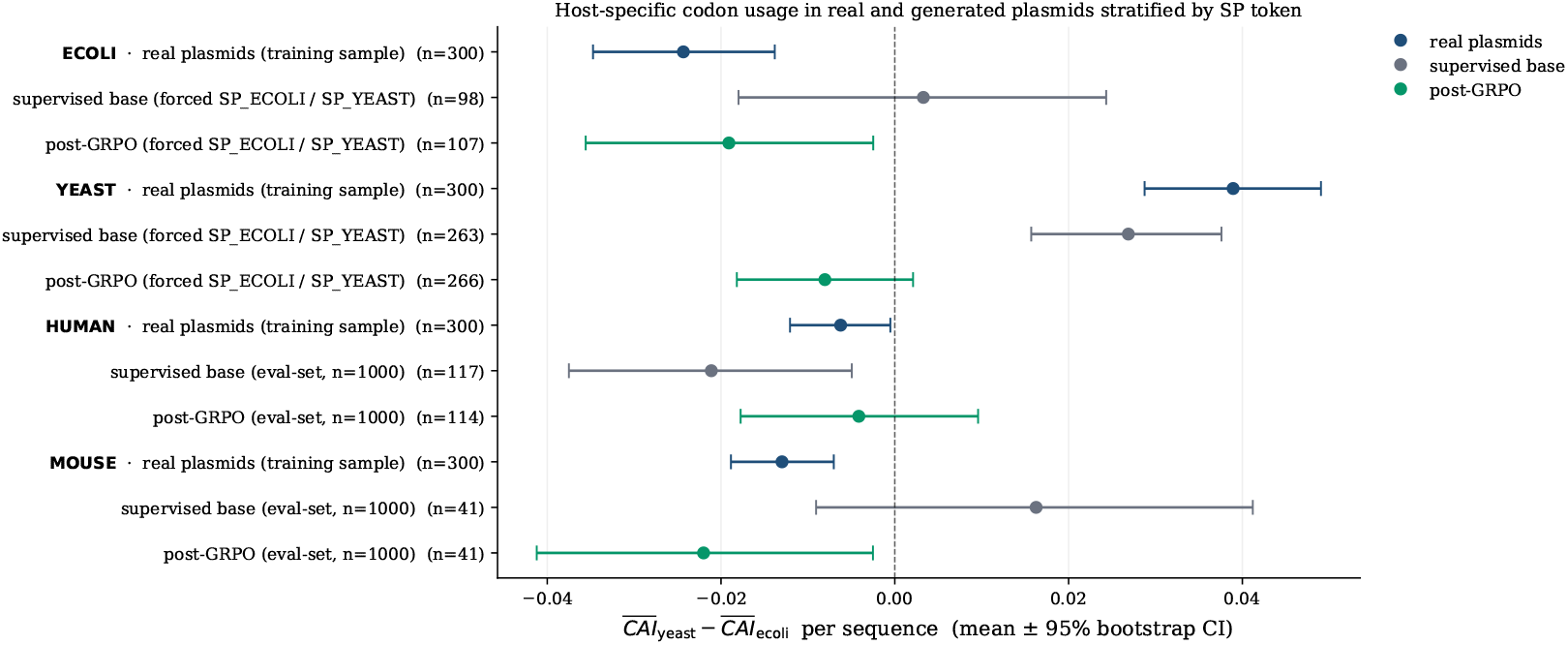
Host-conditional codon usage. For each plasmid we compute ∆ = CAI_yeast_ −CAI_ecoli_ across predicted ORFs (Prodigal). The yeast-vs-*E. coli* contrast isolates codon-preference signal from the GC confound that affects Sharp–Li CAI. Real plasmids show a clean diagonal across four host classes (top section). Generated plasmids reproduce the diagonal partially: the post-GRPO model direction-matches real plasmids on three of four host classes; the supervised base on two of four. Bars are bootstrap 95% CIs over plasmids; rows whose CI excludes zero are significant at *p <* 0.05.

## Notes

### Competing Interest Statement

The authors have declared no competing interest.

